# Machine learning classification by fitting amplicon sequences to existing OTUs

**DOI:** 10.1101/2022.09.01.506299

**Authors:** Courtney R. Armour, Kelly L. Sovacool, William L. Close, Begüm D. Topçuoğlu, Jenna Wiens, Patrick D. Schloss

## Abstract

The ability to use 16S rRNA gene sequence data to train machine learning classification models offers the opportunity diagnose patients based on the composition of their microbiome. In some applications the taxonomic resolution that provides the best models may require the use of *de novo* OTUs whose composition changes when new data are added. We previously developed a new reference-based approach, OptiFit, that fits new sequence data to existing *de novo* OTUs without changing the composition of the original OTUs. While OptiFit produces OTUs that are as high quality as *de novo* OTUs, it is unclear whether this method for fitting new sequence data into existing OTUs will impact the performance of classification models relative to models trained and tested only using *de novo* OTUs. We used OptiFit to cluster sequences into existing OTUs and evaluated model performance in classifying a dataset containing samples from patients with and without colonic screen relevant neoplasia (SRN). We compared the performance of this model to standard methods including *de novo* and database-reference-based clustering. We found that using OptiFit performed as well or better in classifying SRNs. OptiFit can streamline the process of classifying new samples by avoiding the need to retrain models using reclustered sequences.

## Importance

There is great potential for using microbiome data to aid in diagnosis. A challenge with *de novo* OTU-based classification models is that 16S rRNA gene sequences are often assigned to OTUs based on similarity to other sequences in the dataset. If data are generated from new patients, the old and new sequences must be reclustered to OTUs and the classification model retrained. Yet there is a desire to have a single, validated model that can be widely deployed. To overcome this obstacle, we applied the OptiFit clustering algorithm to fit new sequence data to existing OTUs allowing for reuse of the model. A random forest model implemented using OptiFit performed as well as the traditional reassign and retrain approach. This result shows that it is possible to train and apply machine learning models based on OTU relative abundance data that do not require retraining or the use of a reference database.

There is increasing interest in training machine learning models to diagnose diseases such as Crohn’s disease and colorectal cancer using the relative abundance of clusters of similar 16S rRNA gene sequences (1, 2). These models have been used to identify sequence clusters that are important for distinguishing between individuals from different disease categories (3). There is also an opportunity to train models and apply them to classify samples from new individuals. For example, a model for colorectal cancer could be trained, “locked down”, and applied to samples from new patients.

To apply these models to new samples the composition of the clusters would need to be independent of the new data. For example, amplicon sequence variants (ASVs) are defined without consideration of sequences in other samples, phylotypes are defined by clustering sequences that have the same taxonomy (e.g., to the same family) when classified using a taxonomy database, and closed reference operational taxonomic units (OTUs) are defined by mapping sequences to a collection of reference OTUs. In contrast, *de novo* approaches cluster sequences based on their similarity to other sequences in the dataset and can change when new data are added. Although it would be preferrable to select an approach that generates stable clusters, there may be cases where OTUs generated by a *de novo* approach outperform those of the other taxonomic levels. In fact, we recently trained machine learning models for classifying patients with and without screen relevant neoplasias (SRNs) in their colons and found that OTUs generated *de novo* using the OptiClust algorithm performed better than those generated using ASVs or at higher taxonomic levels (4).

It could be possible to construct reference OTUs and map new sequences to those OTUs to attain similar performance as was seen with the OptiClust-generated OTUs. The traditional approach to reference-based clustering of sequences to OTUs has multiple drawbacks and does not produce clusters as good as those generated using OptiClust (5). Sovacool *et al* recently described OptiFit, a method for fitting new sequence data into existing OTUs that overcomes the limitations of traditional reference-based clustering (5). OptiFit allows researchers to fit new data into existing OTUs defined from the same dataset resulting in clusters that are as good as if they had all been clustered with OptiClust. We tested whether OptiClust-generated OTUs could be used to train models that were then used to classify held out samples after clustering their sequences to the model’s OTUs using OptiFit.

To test how the model performance compared between using *de novo* and reference-based clustering approaches, we used a publicly available dataset of 16S rRNA gene sequences from stool samples of healthy subjects (n = 226) as well as subjects with screen-relevant neoplasia (SRN) consisting of advanced adenoma and carcinoma (n = 229) (1). For the *de novo* workflows, the 16S rRNA sequence data from all samples were clustered into OTUs using the OptiClust algorithm in mothur 1 and the VSEARCH algorithm used in QIIME2 (6, 7). For both algorithms, the resulting abundance data was then split into training and testing sets, where the training set was used to tune hyperparameters and ultimately train and select the model. The model was applied to the testing set and performance evaluated (Figure 1A). For traditional reference-based clustering (database-reference-based), we used OptiFit to fit the sequence data into OTUs based on the commonly used greengenes reference database. To compare with another commonly used method, we also used VSEARCH to map sequences to reference OTUs from the greengenes database with the parameters used by QIIME2. Again, the data was then split into training and testing sets, hyperparameters tuned, and performance evaluated on the testing set (Figure 1B). In the OptiFit self-reference workflow (self-reference-based), the data was split into a training and a testing set. The training set was clustered into OTUs and used to train a classification model. The OptiFit algorithm was used to fit sequence data of samples not part of the training data into the training OTUs and classified using the best hyperparameters (Figure 1C). For each of the workflows the process was repeated for 100 random splits of the data to account for variation caused by the choice of the random number generator seed.

**Figure 1:**
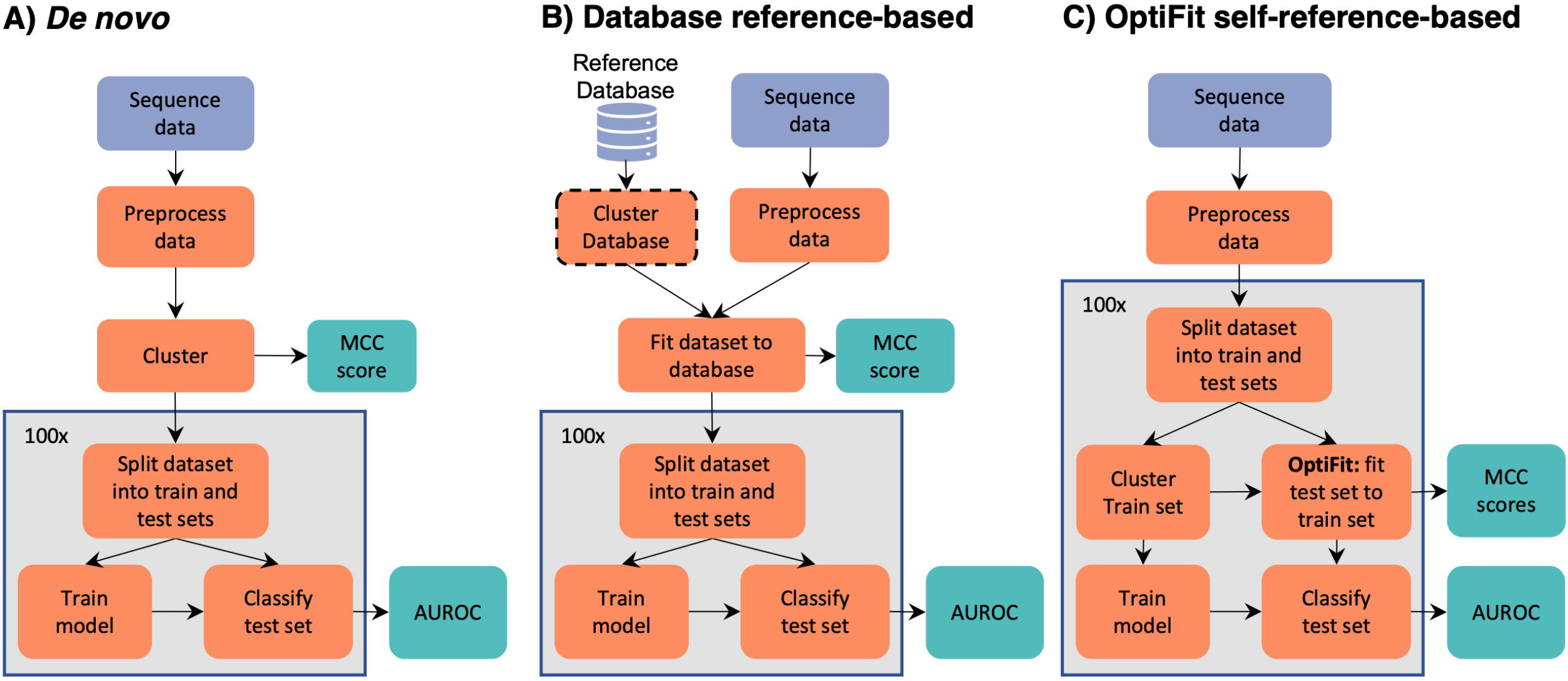
Overview of clustering workflows. The *de novo* and database-reference-based workflows were conducted using two approaches: OptiClust with mothur and VSEARCH as is used in the QIIME pipeline.

We first examined the quality of the resulting OTU clusters from each method using the Matthews correlation coefficient (MCC). MCC is an objective metric used to measure OTU cluster quality based on the similarity of all pairs of sequences and whether they are appropriately clustered or not (8). We expected the MCC scores produced by the OptiFit workflow to be similar to that of *de novo* clustering using the OptiClust algorithm. In the OptiFit workflow the test data was fit to the clustered training data for each of the 100 data splits resulting in an MCC score for each split of the data. In the remaining workflows, the data was only clustered once and then split into the training and testing sets resulting in a single MCC score for each method. Indeed, the MCC scores were similar between the OptiClust *de novo* (MCC = 0.884) and OptiFit self-reference workflows (average MCC = 0.879, standard deviation = 0.002). Consistent with prior findings, the reference-based methods produced lower MCC scores (OptiFit greengenes MCC = 0.786; VSEARCH greengenes MCC = 0.531) than the *de novo* methods (OptiClust *de novo* MCC = 0.884; VSEARCH *de novo* MCC = 0.641) (5). Another metric we examined for the OptiFit workflow was the fraction of sequences from the test set that mapped to the reference OTUs. Since sequences that did not map to reference OTUs were eliminated, if a high percentage of reads did not map to an OTU we expected this loss of data to negatively impact classification performance. We found that loss of data was not an issue since on average 99.8% (standard deviation = 0.7%) of sequences in the subsampled test set mapped to the reference OTUs. This number is higher than the average fraction of reads mapped in the OptiFit greengenes workflow (mean = 96.8%, standard deviation = 3.5). These results indicate that the OptiFit self-reference method performed as well as the OptiClust *de novo* method and better than using an external database.

We next assessed model performance using OTU relative abundances from the training data from the workflows to train a model to predict SRNs and used the model on the held-out data. Using the predicted and actual diagnosis classification, we calculated the area under the receiver operating characteristic curve (AUROC) for each data split. During cross-validation (CV) training, the performance of the OptiFit self-reference and OptiClust *de novo* models were not significantly different (p-value = 0.066; Figure 2A), while performance for both VSEARCH methods was significantly lower than the OptiClust *de novo*, OptiFit self, and OptiFit greengenes methods (p-values < 0.05). The trained model was then applied to the test data classifying samples as either control or SRN. The VSEARCH greengenes method performed slightly worse than the OptiClust *de novo* method (p-value = 0.030). However, the performance on the test data for the OptiClust *de novo*, OptiFit greengenes, OptiFit self-reference, and VSEARCH *de novo* approaches were not significantly different (p-values > 0.05; Figures 2B and 2C). These results indicate that new data could be fit to existing OTU clusters using OptiFit without impacting model performance.

**Figure 2:**
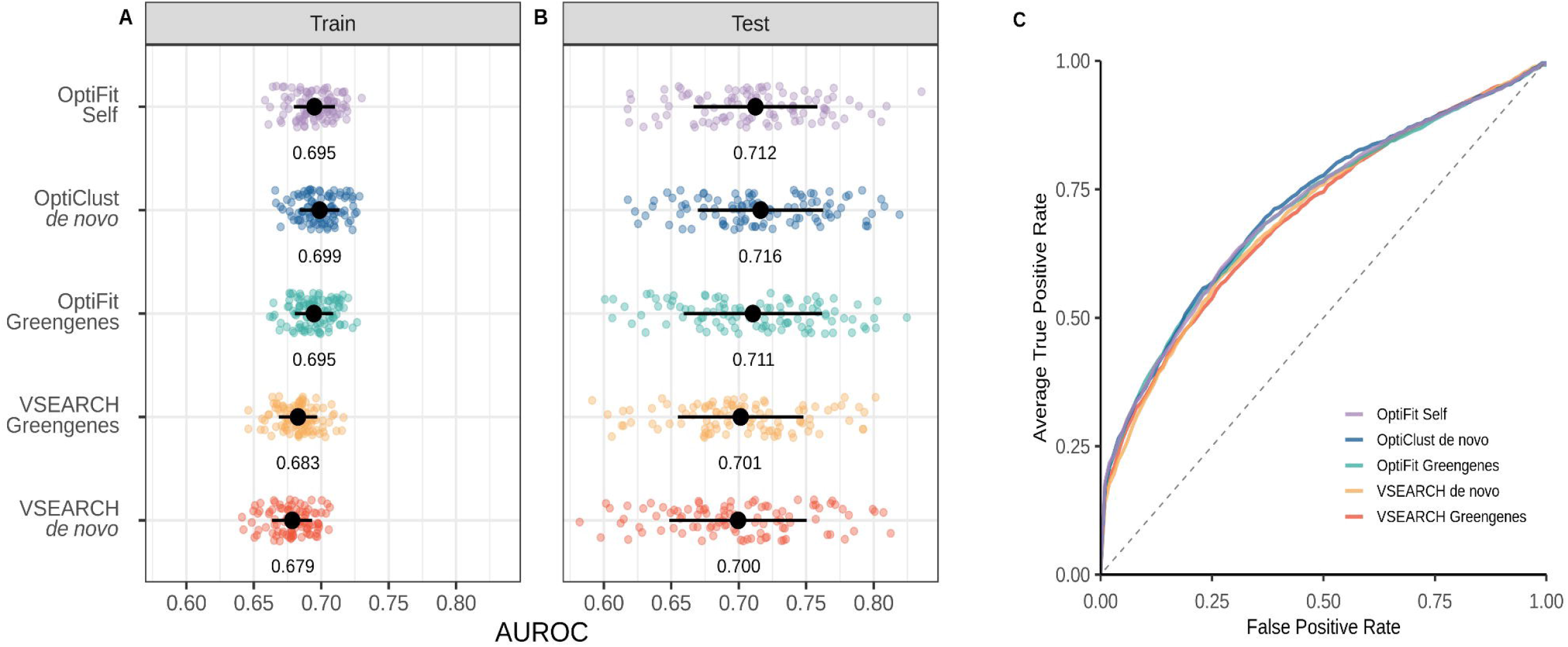
Model performance of OptiFit self-reference workflow is as good or better than other methods. **A)** Area under the receiver operating characteristic (AUROC) curve during cross-validation (train) for the various workflows. **B)** AUROC on the test data for the various workflows. The mean and standard deviation of the AUROC is represented by the black dot and whiskers in panels A and B. The mean AUROC is printed below the points. **C)** Averaged receiver operating characteristic (ROC) curves. Lines represent the average true positive rate for the range of false positive rates.

Random forest machine learning models trained using OptiClust-generated OTUs and tested using OptiFit-generated OTUs performed as well as a model trained using entirely *de novo* OTU assignments. A potential problem with reference-based clustering methods is that sequences that do not map to the reference OTUs are discarded, resulting in a possible loss of information. However, we demonstrated that the training samples represented the most important OTUs for classifying samples. Missing important OTUs is more of a risk when using a database-reference-based method since not all environments are well represented in public databases. Despite this and the lower quality OTUs, the database-reference-based approach performed as well as the models generated using OptiFit. This likely indicates that the sequences that were important to the model were well characterized by the greengenes reference OTUs. However, a less well studied system may not be as well characterized by a reference-database which would make the ability to utilize one’s own data a reference an exciting possibility. Our results highlight that OptiFit overcomes a significant limitation with machine learning models trained using *de novo* OTUs. This is an important result for those applications where models trained using *de novo* OTUs outperform models generated using methods that produce clusters that do not depend on which sequences are included in the dataset.

## Materials and Methods

### Dataset

Raw 16S rRNA gene sequence data from the V4 region were previously generated from human stool samples. Sequences were downloaded from the NCBI Sequence Read Archive (accession no. SRP062005) (1). This dataset contains stool samples from 490 subjects. For this analysis, samples from subjects identified in the metadata as normal, high risk normal, or adenoma were categorized as “normal”, while samples from subjects identified as advanced adenoma or carcinoma were categorized as “screen relevant neoplasia” (SRN). The resulting dataset consisted of 261 normal samples and 229 SRN samples.

### Data processing

The full dataset was preprocessed with mothur (v1.47) (9) to join forward and reverse reads, merge duplicate reads, align to the SILVA reference database (v132) (10), precluster, remove chimeras with UCHIME (11), assign taxonomy, and remove non-bacterial reads following the Schloss Lab MiSeq standard operating procedure described on the mothur website (https://mothur.org/wiki/miseq_sop/). 100 splits of the 490 samples were generated where 80% of the samples (392 samples) were randomly assigned to the training set and the remaining 20% (98 samples) were assigned to the test set. Using 100 splits of the data accounts for the variation that may be observed depending on the samples that are in the training or test sets. Each sample was in the training set an average of 80 times (standard deviation = 4.1) and the test set an average of 20 times (standard deviation = 4.1).

### Reference-based workflows

1. OptiFit Self: The preprocess data was split into the training and testing sets. The training set was clustered into OTUs using OptiClust, then the test set was fit to the OTUs of the training set using the OptiFit algorithm (5). The OptiFit algorithm was run with method open so that any sequences that did not map to the existing OTU clusters would form new OTUs. The data was then subsampled to 10,000 reads and any novel OTUs from the test set were removed. This process was repeated for each of the 100 splits resulting in 100 training and testing datasets.
2. OptiFit Greengenes: Reference sequences from the Greengenes database v13_8_99 (12) were downloaded and processed with mothur by trimming to the V4 region and clustered *de novo* with OptiClust
  a. The preprocessed data was fit to the clustered reference data using OptiFit with the method open to allow any sequences that did not map to the existing reference clusters would form new OTUs. The data was then subsampled to 10,000 reads and any novel OTUs from the test set were removed. The dataset was then split into two sets where 80% of the samples were assigned to the training set and 20% to the testing set. This process was repeated for each of the 100 splits resulting in 100 training and testing datasets.
3. VSEARCH Greengenes: Preprocessed data was clustered using VSEARCH v2.15.2 (6) directly to unprocessed Greengenes 97% OTU reference alignment consistent with how VSEARCH is typically used by the QIIME2 software for reference-based clustering (7). The data was then subsampled to 10,000 reads and any novel OTUs from the test set were removed. The dataset was then split into two sets where 80% of the samples were assigned to the training set and 20% to the testing set. This process was repeated for each of the 100 splits resulting in 100 training and testing datasets.

### De novo workflows

1. OptiClust *de novo*: All the preprocessed data was clustered together with OptiClust 1 to generate OTUs. The data was subsampled to 10,000 reads per sample and the resulting abundance tables were split into the training and testing sets. The process was repeated for each of the 100 splits resulting in 100 training and testing datasets.
2. VSEARCH *de novo*: All the preprocessed data was clustered using VSEARCH v2.15.2 (6) with 97% identity and then subsampled to 10,000 reads per sample. The process was repeated for each of the 100 splits resulting in 100 training and testing datasets for both workflows.

### Machine Learning

A random forest model was trained with the R package mikrompl (v 1.2.0) (13) to predict the diagnosis (SRN or normal) for the samples in the test set for each data split. The training set was preprocessed to normalize OTU counts (scale and center), collapse correlated OTUs, and remove OTUs with zero variance. The preprocessing from the training set was then applied to the test set. Any OTUs in the test set that were not in the training set were removed. P-values comparing model performance were calculated as previously described (14). The averaged ROC curves were plotted by taking the average and standard deviation of the sensitivity at each specificity value.

### Code Availability

The analysis workflow was implemented in Snakemake (15). Scripts for analysis were written in R (16) and GNU bash (17). The software used includes mothur v1.47.0 (9), VSEARCH v2.15.2 (6), RStudio (18), the Tidyverse metapackage (19), R Markdown (20), the SRA toolkit (21), and conda (22). The complete workflow and supporting files required to reproduce this study are available at: https://github.com/SchlossLab/Armour_OptiFitGLNE_XXXX_2023

## Acknowledgments

This work was supported through a grant from the NIH (R01CA215574).

## References

1. Baxter NT, Ruffin MT, Rogers MAM, Schloss PD. 2016. Microbiota-based model improves the sensitivity of fecal immunochemical test for detecting colonic lesions. Genome Medicine 8:37. doi:10.1186/s13073-016-0290-3.

2. Gevers D, Kugathasan S, Denson LA, Vázquez-Baeza Y, Van Treuren W, Ren B, Schwager E, Knights D, Song SJ, Yassour M, Morgan XC, Kostic AD, Luo C, González A, McDonald D, Haberman Y, Walters T, Baker S, Rosh J, Stephens M, Heyman M, Markowitz J, Baldassano R, Griffiths A, Sylvester F, Mack D, Kim S, Crandall W, Hyams J, Huttenhower C, Knight R, Xavier RJ. 2014. The treatment-naive microbiome in new-onset crohn’s disease. Cell Host & Microbe 15:382–392. doi:10.1016/j.chom.2014.02.005.

3. Duvallet C, Gibbons SM, Gurry T, Irizarry RA, Alm EJ. 2017. Meta-analysis of gut microbiome studies identifies disease-specific and shared responses. Nature Communications 8:1784. doi:10.1038/s41467-017-01973-8.

4. Armour CR, Topçuoğlu BD, Garretto A, Schloss PD. 2022. A goldilocks principle for the gut microbiome: Taxonomic resolution matters for microbiome-based classification of colorectal cancer. MBio 13:e0316121.

5. Sovacool KL, Westcott SL, Mumphrey MB, Dotson GA, Schloss PD. 2022. OptiFit: An improved method for fitting amplicon sequences to existing OTUs. mSphere 7:e00916–21. doi:10.1128/msphere.00916-21.

6. Rognes T, Flouri T, Nichols B, Quince C, Mahé F. 2016. VSEARCH: a versatile open source tool for metagenomics. PeerJ 4:e2584. doi:10.7717/peerj.2584.

7. Bolyen E, Rideout JR, Dillon MR, Bokulich NA, Abnet CC, Al-Ghalith GA, Alexander H, Alm EJ, Arumugam M, Asnicar F, Bai Y, Bisanz JE, Bittinger K, Brejnrod A, Brislawn CJ, Brown CT, Callahan BJ, Caraballo-Rodríguez AM, Chase J, Cope EK, Da Silva R, Diener C, Dorrestein PC, Douglas GM, Durall DM, Duvallet C, Edwardson CF, Ernst M, Estaki M, Fouquier J, Gauglitz JM, Gibbons SM, Gibson DL, Gonzalez A, Gorlick K, Guo J, Hillmann B, Holmes S, Holste H, Huttenhower C, Huttley GA, Janssen S, Jarmusch AK, Jiang L, Kaehler BD, Kang KB, Keefe CR, Keim P, Kelley ST, Knights D, Koester I, Kosciolek T, Kreps J, Langille MGI, Lee J, Ley R, Liu Y-X, Loftfield E, Lozupone C, Maher M, Marotz C, Martin BD, McDonald D, McIver LJ, Melnik AV, Metcalf JL, Morgan SC, Morton JT, Naimey AT, Navas-Molina JA, Nothias LF, Orchanian SB, Pearson T, Peoples SL, Petras D, Preuss ML, Pruesse E, Rasmussen LB, Rivers A, Robeson MS, Rosenthal P, Segata N, Shaffer M, Shiffer A, Sinha R, Song SJ, Spear JR, Swafford AD, Thompson LR, Torres PJ, Trinh P, Tripathi A, Turnbaugh PJ, Ul-Hasan S, Hooft JJJ van der, Vargas F, Vázquez-Baeza Y, Vogtmann E, Hippel M von, Walters W, Wan Y, Wang M, Warren J, Weber KC, Williamson CHD, Willis AD, Xu ZZ, Zaneveld JR, Zhang Y, Zhu Q, Knight R, Caporaso JG. 2019. Reproducible, interactive, scalable and extensible microbiome data science using QIIME 2. Nature Biotechnology 37:852–857. doi:10.1038/s41587-019-0209-9.

8. Westcott SL, Schloss PD. 2015. De novo clustering methods outperform reference-based methods for assigning 16S rRNA gene sequences to operational taxonomic units. PeerJ 3:e1487. doi:10.7717/peerj.1487.

9. Schloss PD, Westcott SL, Ryabin T, Hall JR, Hartmann M, Hollister EB, Lesniewski RA, Oakley BB, Parks DH, Robinson CJ, Sahl JW, Stres B, Thallinger GG, Van Horn DJ, Weber CF. 2009. Introducing mothur: Open-source, platform-independent, community-supported software for describing and comparing microbial communities. Applied and Environmental Microbiology 75:7537–7541. doi:10.1128/AEM.01541-09.

10. Quast C, Pruesse E, Yilmaz P, Gerken J, Schweer T, Yarza P, Peplies J, Glöckner FO. 2013. The SILVA ribosomal RNA gene database project: Improved data processing and web-based tools. Nucleic Acids Research 41:D590–D596. doi:10.1093/nar/gks1219.

11. Edgar RC, Haas BJ, Clemente JC, Quince C, Knight R. 2011. UCHIME improves sensitivity and speed of chimera detection. Bioinformatics 27:2194–2200. doi:10.1093/bioinformatics/btr381.

12. DeSantis TZ, Hugenholtz P, Larsen N, Rojas M, Brodie EL, Keller K, Huber T, Dalevi D, Hu P, Andersen GL. 2006. Greengenes, a chimera-checked 16S rRNA gene database and workbench compatible with ARB. Applied and Environmental Microbiology 72:5069–5072. doi:10.1128/AEM.03006-05.

13. Topçuoğlu BD, Lapp Z, Sovacool KL, Snitkin E, Wiens J, Schloss PD. 2021. mikropml: User-Friendly R Package for Supervised Machine Learning Pipelines. Journal of Open Source Software 6:3073. doi:10.21105/joss.03073.

14. Topçuoğlu BD, Lesniak NA, Ruffin MT, Wiens J, Schloss PD. 2020. A framework for effective application of machine learning to microbiome-based classification problems. mBio 11:e00434–20. doi:10.1128/mBio.00434-20.

15. Koster J, Rahmann S. 2012. Snakemake–a scalable bioinformatics workflow engine. Bioinformatics 28:2520–2522. doi:10.1093/bioinformatics/bts480.

16. R Core Team. 2020. R: A language and environment for statistical computing. R Foundation for Statistical Computing, Vienna, Austria.

17. GNU Project. Bash reference manual.

18. RStudio Team. 2019. RStudio: Integrated development environment for r. RStudio, Inc., Boston, MA.

19. Wickham H, Averick M, Bryan J, Chang W, McGowan LD, François R, Grolemund G, Hayes A, Henry L, Hester J, Kuhn M, Pedersen TL, Miller E, Bache SM, Müller K, Ooms J, Robinson D, Seidel DP, Spinu V, Takahashi K, Vaughan D, Wilke C, Woo K, Yutani H. 2019. Welcome to the Tidyverse. Journal of Open Source Software 4:1686. doi:10.21105/joss.01686.

20. Xie Y, Allaire JJ, Grolemund G. 2018. R Markdown: The Definitive Guide. Taylor & Francis, CRC Press.

21. SRA-Tools - NCBI. http://ncbi.github.io/sra-tools/.

22. 2016. Anaconda Software Distribution. Anaconda Documentation. Anaconda Inc.

